# Investigation of autophagy-activating molecules in a glia-specific Spinocerebellar ataxia type 1 model

**DOI:** 10.64898/2026.02.23.707351

**Authors:** Tímea Burján, Hanna Horváth, Eszter Illés, Katalin Schlett, Norbert Bencsik, Tibor Kovács

## Abstract

Autophagy is a critical neuroprotective mechanism, the impairment of which can lead to severe neurodegenerative diseases. Spinocerebellar ataxia type 1 (SCA1) is a monogenic neurodegenerative disorder, characterised by the presence of protein aggregates and consequent loss of cellular functions. The expression of mutant Ataxin1 (ATXN1) in glial cells has been demonstrated to induce inflammatory responses and loss of supportive functions, thereby exacerbating neuronal degeneration in SCA1. Autophagic dysfunction has been shown to affect both neurons and glial cells, resulting in widespread pathological consequences. In this work, we aimed to evaluate the efficacy of two small-molecule autophagy activators, AUTEN-67 and AUTEN-99, in models of glia-specific SCA1 in *Drosophila*. Our results demonstrate that AUTEN-99 has a stronger autophagy enhancing effect, with significantly improved response times and survival rates, compared to untreated ATXN1 mutants. Glia-specific assays in mouse primary hippocampal cultures also confirmed that AUTEN-99 is a more effective activator. Ultimately, co-treatment of neuronal and glial cultures did not reveal any synergistic benefits from combining the two AUTEN compounds compared to single-agent treatment. Our findings contribute to a better understanding of the utility of AUTENs and may help to understand the critical role of autophagy in neurodegenerative diseases.

## Introduction

Spinocerebellar ataxia type 1 (SCA1) is an incurable, autosomal dominant monogenic neurodegenerative disorder caused by pathological CAG repeat expansion in the ATXN1 gene ^1,2^. Clinically, SCA1 is a progressive disorder predominantly affecting the cerebellum and brainstem, manifesting with ataxia, loss of balance, dysarthria, dysphagia and oculomotor abnormalities ^3^. Mutant ATXN1 protein accumulates in toxic nuclear and cytoplasmic aggregates within Purkinje neurons, leading to synaptic dysfunction, dendritic atrophy, impaired firing and eventual cell death — changes that underlie the ataxic phenotype ^2,4^. Recent studies indicate that SCA1 associated cellular alterations can be detected in glial cells prior to overt neuronal changes ^5^. Expression of mutant ATXN1 in glia, notably cerebellar Bergmann glia, induces inflammatory signalling, loss of supportive functions and disruption of neuron–glia homeostasis, thereby non-cell-autonomously exacerbating neuronal dysfunction and degeneration ^6^. Astrocytes play neuroprotective roles in early disease stages but adopt a chronically reactive, proinflammatory state in later phases, thereby potentiating neuronal loss ^7^. Oligodendrocytes show transcriptional perturbations before the appearance of overt motor symptoms, and their dysfunction reduces the efficiency of axonal impulse conduction ^5^. SCA1 is a dominant monogenic neurodegenerative disorder caused by pathological CAG repeat expansion in the ATXN1 gene ^8^. Associated cellular alterations can be detected in glial cells prior to overt neuronal changes, as these cells non-autonomously exacerbate neuronal dysfunction and degeneration, and an inflammatory state in later phases, thereby potentiating neuronal loss ^9,10^.

Autophagy is a conserved intracellular degradative pathway that delivers cytoplasmic cargo — damaged organelles, protein aggregates and pathogens — to lysosomes for breakdown and recycling, thereby maintaining proteostasis and cellular homeostasis ^11–13^. Macroautophagy (hereafter autophagy) is the principal route by which double-membraned autophagosomes sequester cytoplasmic material and subsequently fuse with lysosomes (forming autolysosomes) for degradation ^14^. Autophagy is regulated by a cascade of complexes acting in series. A key step in autophagosome biogenesis is vesicle nucleation, during which membranes of various intracellular origins are designated for autophagic processing. This designation is executed by the Vps34 kinase complex, which phosphorylates phosphatidylinositol (PI) to produce phosphatidylinositol-3-phosphate (PI3P), a lipid that recruits downstream autophagy machinery ^15–17^. Antagonists of this phosphorylation include myotubularin family phosphatases (MTMRs), notably MTMR14. MTMR14 (Jumpy) and its *Drosophila* orthologue EDTP (Egg-derived tyrosine phosphatase) dephosphorylate PI3P back to PI, thereby inhibiting autophagy ^18^. In the nervous system, autophagic capacity declines with advancing age, whereas MTMR14/EDTP expression increases, promoting progressive inhibition of lysosomal degradation ^19,20^. Genetic inhibition of EDTP has been reported to extend lifespan and improve cognitive function ^20^. Numerous autophagy-inducing agents can be used to stimulate autophagy in ageing. However, many of these — including the well-known rapamycin — act upstream of autophagy initiation and therefore affect multiple cellular pathways ^21^. The Vps34 vesicle-nucleation complex is a more downstream regulatory node; activating it or inhibiting its antagonists offers a more specific means of modulating autophagosome biogenesis ^22^.

Two pharmacological inhibitors of MTMR14 have been described: AUTEN67 and AUTEN99 ^23,24^. AUTEN treatments have been shown to extend lifespan in invertebrate models. In multiple neurodegenerative tests (Parkinson’s, Alzheimer’s, and Huntington’s disease models), AUTENs reduced the levels of toxic proteins and aggregates, extended expected lifespan and ameliorated disease-relevant phenotypes; in a *Drosophila* Parkinson’s model, AUTEN treatment reduced dopaminergic neuron loss ^24,25^. In our recent study, we compared the actions of AUTEN67 and AUTEN99 across different neuronal types and observed differential sensitivity of distinct neuronal subpopulations to these compounds ^26^. The same study evaluated the sensitivity of neurons expressing mutant ATXN1 in a cell-type-specific manner to AUTENs, but did not assess the separate or combined effects of AUTEN67 and AUTEN99 on glial cells ^26^.

## Results

### AUTEN-99 induces autophagy in Drosophila glial cells

Our goal was to study the impact of the small molecules AUTEN-67 and AUTEN-99 on autophagy in glial cells. We established an experimental system that allows us to assess autophagy exclusively in glial cells. Using the glia-specific repo-Gal4 driver, we expressed the UAS-GFP-mCherry-Atg8a marker in Drosophila. Following seven days of AUTEN treatment (both were used at a concentration of 100 µM), we analyzed glial cells in the adult animals’ brains ^27^. Atg8a, the Drosophila ortholog of LC3B, covalently binds to the forming phagophore membrane and remains detectable in all autophagic vesicles up to the autolysosome stage ^28^. GFP is pH-sensitive and rapidly loses its fluorescence in acidic environments such as those in autophagosomes fused with late endosomes or lysosomes. In contrast, the mCherry signal remains stable and detectable even at low pH. Consequently, a defect of the fusion results in the accumulation of yellow Atg8a-positive vesicles (GFP and mCherry colocalization), while an increase in red-only Atg8a-positive vesicles (mCherry alone) indicates the accumulation of amphisomes (vesicles formed by the fusion of autophagosomes with late endosomes) and autolysosomes ^29^. Since additional autophagosomes can fuse with pre-existing autolysosomes, the size of autolysosomes becomes a more informative parameter over time than the number of the structures ^20^. Therefore, we quantified changes in vesicle size in our measurements. Since the autophagy-activating compounds were dissolved in dimethyl sulfoxide (DMSO), the untreated control group received the same concentration of DMSO as the AUTEN-treated groups.

Analysis of the three experimental groups revealed no accumulation of GFP-mCherry double-positive vesicles under any conditions. This indicates that fusion of autophagosomes with acidic vesicles was not impaired in any of the samples (**Figure 1A**). The mCherry-Atg8a signal showed a significant increase in the abundance of autophagic structures in AUTEN-99-treated samples (**Figure 1B, B’**). Proteins containing FYVE domains can bind to phosphatidylinositol-3-phosphate (PI3P), which provides information about the activity of the Vps34 vesicle nucleation complex ^30,31^. Myotubularin-related protein 14 (MTMR14) and its Drosophila ortholog, egg-derived tyrosine phosphatase (EDTP), act as antagonists of the Vps34 complex by dephosphorylating PI3P to phosphatidylinositol (PI) ^18,32^. The efficacy of AUTEN-67 and AUTEN-99 is based on their inhibitory effects on MTMR14 and EDTP activity.^23,24^. Using the GFP-2xFYVE marker, we assessed cellular PI3P levels ^33^. We found that PI3P levels increased with treatment by both AUTEN compounds, though this increase was only statistically significant in AUTEN-99-treated samples (**Figure 1C,C’**). Finally, we examined the efficiency of autophagic degradation (autophagic flux) in AUTEN-treated glial cells. For this analysis, we isolated proteins from the heads of repo-Gal4-driven GFP-mCherry-Atg8a-expressing flies. Western blot analysis using an anti-GFP antibody revealed four distinct bands. The upper bands correspond to the soluble GFP-mCherry-Atg8a-I and lipidated GFP-mCherry-Atg8a-II forms. The free mCherry-GFP and GFP proteins represent C-terminal cleavage products generated by acidic hydrolases during the degradation of Atg8a by autophagy ^26,34^. Our results demonstrate a significant increase in cleaved GFP and mCherry-GFP fragment levels in AUTEN-99-treated samples (**Figure 1D, D’**). Taken together, these findings suggest that AUTEN-99 significantly enhances autophagy in *Drosophila* glial cells and is more effective than AUTEN-67 at the same concentration.

**Figure 1:**
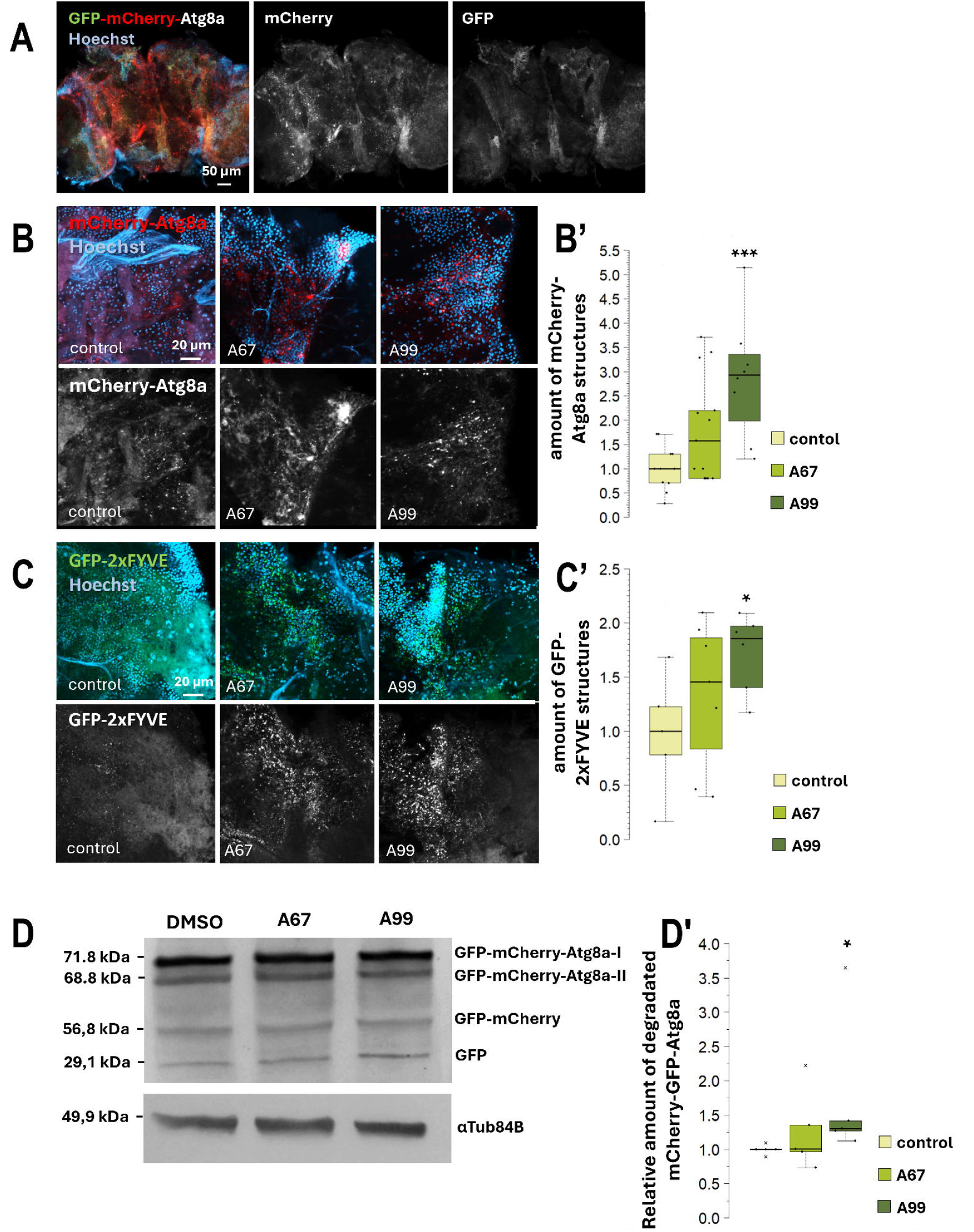
AUTEN-99 increases autophagy in Drosophila glial cells to a greater extent than AUTEN-67. **A)** To assess autophagy activity, GFP-mCherry-Atg8a was expressed from a UAS promoter under repo-Gal4 control (a glial-specific Gal4 driver). The number of GFP puncta did not increase in any sample, indicating that autophagy was not blocked after Atg8a lipidation. Panel A shows repo-Gal4-driven expression in a dissected adult Drosophila brain. B–B′) The number of mCherry-positive vesicles (autolysosomes; red) increased significantly in the glial cells of AUTEN-99-treated brains. C–C′) GFP-2xFYVE (green) binds phosphatidylinositol 3-phosphate (PI3P). MTM14/EDTP is an antagonist of the Vps34 complex; inhibiting it elevates PI3P levels. AUTEN-99 significantly increased the abundance of GFP-2xFYVE structures C’). Nuclei were stained with Hoechst (blue) in the confocal images. D–D′) Protein extracts were prepared from AUTEN-67-, AUTEN-99-treated, and control (DMSO) animals expressing repo-Gal4-driven GFP-mCherry-Atg8a in glia. To determine autophagic flux, cleaved GFP and GFP-mCherry fragments were compared using anti-GFP immunoblotting. AUTEN-99-treated glia exhibited elevated levels of free GFP and GFP-mCherry fragments. αTub84B was used as an internal control. Together, these results suggest that AUTEN-99 increases autophagy in glial cells of the adult Drosophila brain.

### AUTEN-99 enhances autophagic activity in primary mouse glial cells

In our previous study, we examined the response of neurons in mouse embryonic hippocampal primary cultures to AUTEN treatments. In these cultures, glial cells are also present ^26,35^ and can be distinguished from neurons based on well-defined morphological characteristics. Neurons typically exhibit round or pyramidal soma with one prominent axon and multiple branching dendrites forming an extended network. In contrast, glial cells typically exhibit a flattened shape with a relatively large polygonal soma, and they often form a confluent monolayer supporting neuronal networks ^36^ (Figure 2A). Lysosomal-associated membrane protein 1 (LAMP1) is a transmembrane glycoprotein predominantly localised to the lysosomal membrane^37^ and is widely used as a lysosomal marker^38^. During 24 hours of 10 μM AUTEN treatment, both compounds increased the abundance of LAMP1-immunoreactive structures. (**Figure 2B,B’**). p62 serves as a selective substrate for autophagic degradation^39^. Its abundance is inversely correlated with autophagic efficiency: a decrease indicates autophagy activation, whereas accumulation reflects impaired lysosomal degradation^39^. In addition to the DMSO-only control, Bafilomycin A1 was used as a negative control during the treatments. Bafilomycin A1 is an autophagy inhibitor that blocks autophagosome–lysosome fusion^40^. In glial cells treated with Bafilomycin A1, the amount of p62-positive structures was significantly increased, indicating autophagy inhibition. (**Figure 2B and S.Figure 1A**). The two AUTEN treatments had opposite effects on the abundance of p62-positive structures. AUTEN-67, similarly to Bafilomycin A1, increased p62 levels, whereas AUTEN-99 significantly decreased them. (**Figure 2B,B’’**). These findings indicate that AUTEN-99 seems to promote autophagy in primary embryonic mouse glial cells, while AUTEN-67 appears to inhibit it in the same context.

**Figure 2:**
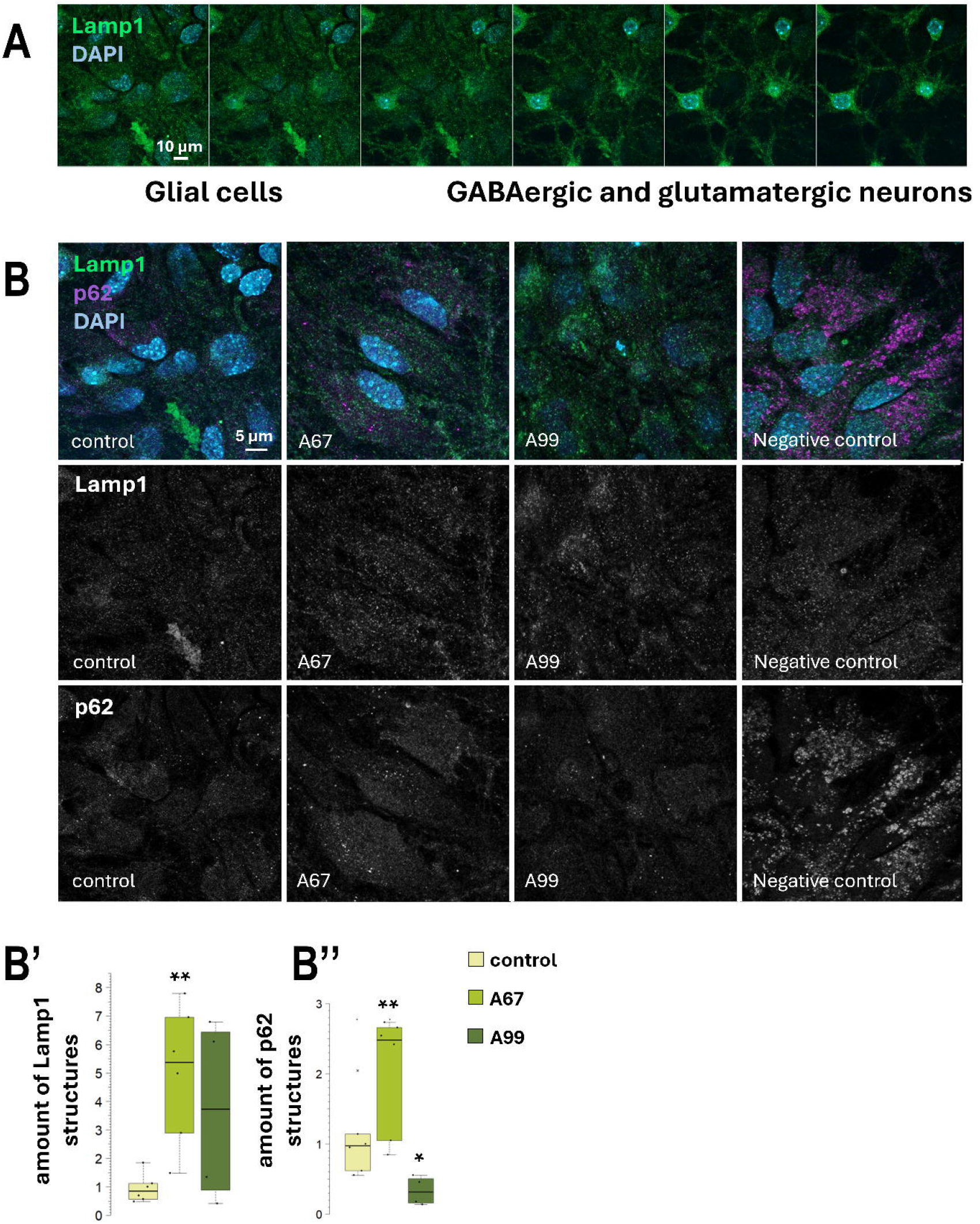
AUTEN-99 increases autophagy in the glial cells of mouse embryonic hippocampal cultures. The treatments used an equivalent volume of dimethyl sulfoxide (DMSO) as the vehicle control (labelled ‘DMSO’ in the panels), which matched the solvent for the two drugs. A) Under basal conditions, mouse primary embryonic hippocampal cultures contain mixed neuronal and glial layers. B–B′) Lysosomal abundance was assessed by LAMP1 immunolabelling following AUTEN-67 (A67) and AUTEN-99 (A99) treatment. Both compounds increased the number of lysosomal structures. B, B′′) p62 (magenta) is an autophagy substrate, the cellular abundance of which is inversely correlated with autophagic efficiency. A67 significantly increased the number of p62-positive structures, whereas A99 reduced them. Overall, these results suggest that A99 enhances autophagy in mammalian glial cells, while A67 likely has an inhibitory effect. Bafilomycin treatment was used as a negative control, which inhibits autophagosome-lysosome fusion, leading to the expected p62 accumulation.

### AUTEN treatment increases survival and responsiveness in Drosophila carrying a glia-specific ATXN1 mutation

To determine whether the differential efficacy of the two AUTEN compounds is also evident in a glia-specific SCA1 model, we overexpressed wild-type (30Q) and mutant (82Q) human ATXN1 in *Drosophila* glial cells using the repo-Gal4 driver. SCA1 is a dominant polyglutamine (polyQ) neurodegenerative disorder, comparable to Huntington’s disease, in which the length of the CAG repeat expansion in the ATXN1 gene is a key determinant of disease development^41^. The 30Q construct represents the non-pathogenic form of ATXN1, while the 82Q construct corresponds to the pathogenic, disease-associated variant^42^. The 82Q repo-Gal4 animals exhibited a pupal lethal phenotype, consistent with the observations reported by Tamura and colleagues^43^. The temperature-sensitive Gal80ts inhibits UAS-Gal4 activity at room temperature, allowing the generation of a model in which 82Q expression is activated only in adults emerging from the pupal stage. Autophagic vesicles were assessed by quantifying punctate mCherry-Atg8a structures. Compared to 30Q controls, the number of mCherry-Atg8a structures was significantly reduced in 82Q control animals. This reduction was significantly rescued by AUTEN-99 treatment, and AUTEN-67 treatment also increased the number of autophagic structures relative to 82Q controls (**Figure 3A,A’**). Lifespan analysis revealed a drastic reduction in animals expressing 82Q in glial cells compared to 30Q controls. This decrease was partially rescued by both AUTEN-99 and AUTEN-67, with AUTEN-99 remaining the more effective of the two compounds (**Figure 3B and S.Figure 1B**). When placed in a narrow, vertical tube, *Drosophilas* exhibit a negative geotaxis escape reflex: they climb upwards after being gently tapped to the bottom of the tube. Since ATXN1 wild-type and mutant proteins were only overexpressed in glial cells, the differences between the two variants can be interpreted independently of locomotor defects. This assay enables the evaluation of the responsiveness of different experimental groups^44^. We found that glial-specific overexpression of the 82Q mutant ATXN1 protein significantly impaired animal responsiveness, which was significantly improved by treatment with both AUTEN compounds (**Figure 3C**).

**Figure 3:**
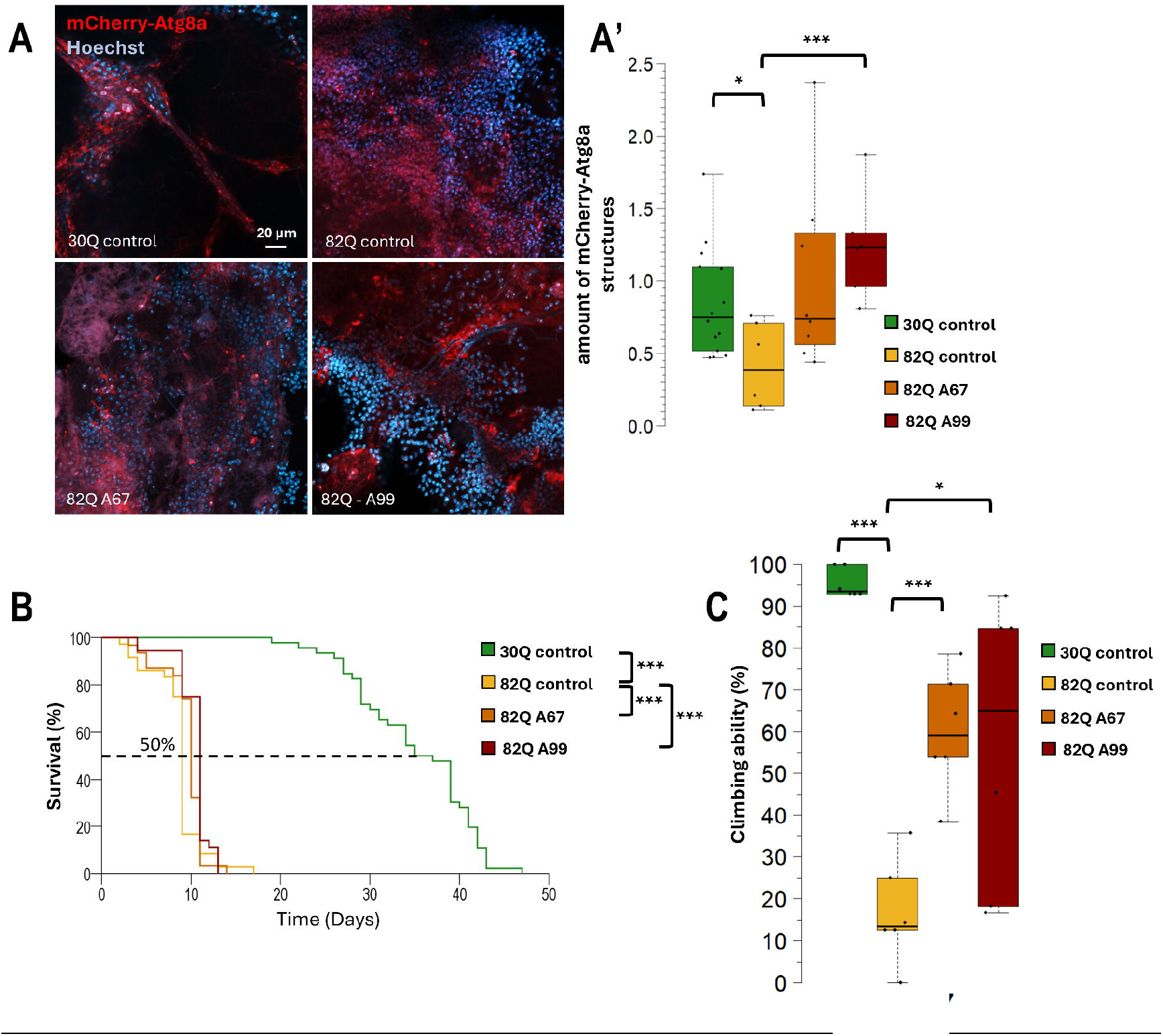
AUTEN-99 is an autophagy activator that increases the lifespan of SCA1 disease models. Wild-type (30Q) and mutant (82Q) ATXN1 proteins were overexpressed in the glia of adult Drosophila using repo-Gal4. Tub-Gal80[ts] was used to restrict UAS-Gal4 activity to adults, thereby avoiding developmental effects. Vehicle controls received an equivalent volume of DMSO to that used to dissolve the AUTENs. A–A’) Autophagic vesicles in glia were quantified using mCherry–Atg8a (repo-Gal4 driven). AUTEN-99 (A99) increased the number of mCherry–Atg8a-positive structures more effectively in 82Q mutant animals than AUTEN-67 (A67) did. B–C) Both A67 and A99 increased the mean survival and behavioural responsiveness of animals expressing 82Q ATXN1 in glia.

## Discussion

Numerous small-molecule autophagy activators have been described; however, the majority act upstream of the Atg1 activation complex. Rapamycin and the Torin1–Torin2 molecules inhibit the TORC1 and TORC2 complexes, which, in addition to promoting autophagy, exert tumour-suppressive effects. Nevertheless, these agents cannot be considered as specific autophagy modulators^45,46^. In addition to suppressing autophagy, TORC1 plays a central role in regulating cell growth, ribosome biogenesis, and protein synthesis. Consequently, inhibiting TORC1 leads to significant and extensive changes in numerous cellular processes^47,48^. Treatment with metformin or 5-Aminoimidazole-4-carboxamide ribonucleotide (AICAR) activates AMP-activated protein kinase (AMPK)^49^. AMPK enhances autophagy via two main mechanisms: by inhibiting TORC1 and by directly activating the Atg1/ULK1 kinase complex^50,51^. For the reasons outlined above, inhibition of TORC1 cannot be regarded as autophagy-specific. New research suggests that AICAR has effects independent of AMPK^52^. Furthermore, Atg1 plays a role in endocytosis in addition to autophagy. Atg1 also exerts negative feedback on TORC1 activity, making it indirectly involved in regulating cell size, protein synthesis, and ribosome biogenesis^53^. BH3 mimetics, such as ABT-737, Navitoclax, and Venetoclax, disrupt the Beclin 1–Bcl-2 complex. This releases Beclin 1 from its inhibitory interaction with Bcl-2^54,55^. The usage of BH3 mimetics usually induces apoptosis by the inhibition of antiapoptotic Bcl-2^56^. Natural compounds can also activate the Vps34 complex. For example, resveratrol induces enhanced AMPK activation, TORC1 inhibition, and subsequent Vps34 activation^57^. Spermidine enhances the activity of the Vps34 complex through inhibition of EP300^58,59^. EP300 is a lysine acetyltransferase that regulates acetylation-dependent processes. Through histone acetylation, it modulates the expression of numerous genes, while acetylation of non-histone proteins influences their stability and subcellular localisation. Inhibition of EP300 alters the acetylation status of several Atg proteins, thereby enhancing autophagy; however, these effects cannot be considered autophagy-specific^59^. Vps34 participates in two distinct complexes that catalyse the phosphorylation of phosphatidylinositol to generate PI3P. The core Vps34–Beclin 1–Vps15 complex associated with Atg14 regulates autophagy, whereas the UVRAG-containing complex primarily controls early endocytic trafficking ^15^.

AUTEN-67 and AUTEN-99 promote autophagy through inhibition of MTMR14, a functional antagonist of the Vps34 complex^23,24^. According to the literature, MTMR14 preferentially reduces the pool of PI3P associated with the autophagic pathway, while exerting only a modest influence on endocytotic PI3P. According to Verge et al., the MTMR14 reporter (GFP-Jumpy) does not colocalize with EEA1-positive endosomal PI3P^60^. The role of EDTP in endocytosis was investigated in UVRAG-deficient Drosophila; in this context, GFP–2xFYVE-positive structures were elevated independently of endocytosis, suggesting that EDTP likewise modulates the autophagic pathway^18^. To date, the effects of AUTEN compounds have been studied in Drosophila and HEK cells. In both models, a minor increase in endocytic structures was observed, while enhanced autophagic activity was noted^23,24^. Based on the current evidence, AUTEN-mediated activation of autophagy may be associated with fewer off-target effects than other autophagy-activating compounds.

Based on our results, glial cells appear to be more sensitive to AUTEN-99 treatment than to AUTEN-67 treatment. This conclusion is supported by the increased presence of mCherry-Atg8a- and GFP-2xFYVE-positive autophagic structures, as well as the increased amount of degraded GFP-mCherry-Atg8a fragments, in Drosophila glia (Figure 1). These findings are consistent with observations in mouse embryonic hippocampal cultures, in which AUTEN-99 increased Lamp1-positive lysosomal structures and reduced p62 levels in treated glial cells (Figure 2). Previously, we compared AUTEN-67 and AUTEN-99 in Drosophila indirect flight muscle (IFM); both compounds reduced p62 aggregate load. In aged flies, AUTEN-treated samples showed a significant decrease in the abnormal mitochondria characteristic of striated muscle. AUTEN treatment mitigated age-associated declines in autophagic activity and preserved flight performance^61^. In contrast, analysis of different neuronal subtypes revealed that the AUTEN compounds had distinct effects on autophagy induction. Both compounds increased autophagy in GABAergic and dopaminergic neurons. However, only AUTEN-67 was effective in cholinergic neurons, and only AUTEN-99 was potent in glutamatergic neurons. In mammalian neuronal cultures, differences between AUTEN-67 and AUTEN-99 were also observed in GABAergic neurons. AUTEN-99 significantly increased Lamp1-positive lysosomal structures and decreased p62-positive structures while AUTEN-67 increased lysosomal structures in GABAergic neurons but did not significantly affect p62 levels^26^.

In glial cells we also observed increases in p62 and Lamp1 following AUTEN-67 treatment (Figure 2), which indicates the inhibition at downstream steps of autophagy^26^. The conversion of PI3P to PI(3,5)P is catalyzed by the PIKfyve complex, which is required for late endosome and lysosome motility and for vesicle–lysosome fusion^62^. Members of the myotubularin-related protein (MTMR) family mediate the dephosphorylation of phosphatidylinositol (PI)(3,5)P_2_ to phosphatidylinositol (PI)5P^63^, hereby inhibiting autophagy. We hypothesized that AUTEN-67 would promote vesicle-lysosome fusion and improve autophagic flux if it acted at this node. Therefore, our results do not support the theory that AUTEN-67 inhibits MTMR14 as a PIKfyve antagonist in glial cells. It is known that catalytically active MTMRs can form complexes with inactive MTMR paralogs, thereby altering their function or substrate specificity ^18,64^. However, MTMR14 lacks the coiled-coil domain required for heterodimerization with other MTMRs^60^. There is no literature evidence identifying another MTMR annotated as forming a complex with MTMR14. An alternative explanation is to question the specificity of the AUTEN compounds for MTMR14: the effects observed with AUTEN-67 could partly reflect inhibition of a different MTMR.

To date, AUTEN specificity has been evaluated in muscle, larval fat body, and the whole Drosophila nervous system. In EDTP mutant animals, neither AUTEN compound was able to further enhance autophagy in those tissues^23,24,61^. We generated glia-specific SCA1 disease models by overexpressing human ATXN1 82Q and 30Q. Both AUTEN compounds increased mCherry-Atg8a–positive structures in glia and improved survival and response time of 82Q animals (**Figure 3**). Across assays AUTEN-99 was consistently slightly more effective than AUTEN-67. Interestingly, in wild-type animals only AUTEN-99 produced a significant activation of autophagy. We hypothesize that AUTEN-67 can elicit a smaller increase in glial autophagy that becomes more apparent under proteotoxic stress (ATXN1 82Q overexpression), when glias are more responsive to induction of cell survival mechanisms.

## Materials and Methods

### Drosophila Strains and AUTEN Treatments

6 *Drosophila* strains were obtained from the Bloomington Drosophila Stock Center (BDSC): w[1118] (BDRC 5905), w[*]; P{w[+mC]=UAS-GFP-myc-2xFYVE}2 (BDRC 42712), P{Appl-GAL4.G1a}1, y1 w* (BDRC 32040), w[*]; P{w[+mC]=ChAT-GAL4.7.4}19B (BDRC 6793), P{w[+mC]=Gad1-GAL4.3.098}2/CyO (BDRC 51630), w[*]; P{w[+mC]=ple-GAL4.F}3 (BDRC 8848). Huda Zoghbi was kind to provide us the following 3 strains: P{w[+mC]=UAS-Hsap\ATX1.30Q}F6 (30Q), w[1118]; P{w[+mC]=UAS-Hsap\ATX1.82Q}M6 (82Q), w[*]; P{w[+mC]=GAL4-elav.L}3 (Elav-Gal4). We also acquired strains from Gábor Juhász’s laboratory: w[*]; P{UAS-mCherry-GFP-Atg8a} (UGMA), w[1118]; 3xmCherry-Atg8a, w[*]; TubGFP-p62 3-2M (II)/CyO, w[*]. And last but not least we obtained flies from Áron Szabó: w[1118]; P{w[+m*]=GAL4}repo/TM3, Sb[1] (BDSC 7415), P{w[+mC]=Gad1-GAL4.3.098}2/CyO (BDSC 51630).

The flies were kept on 25°C and standard medium. Treatments and experiments were performed after hatching from the pupae at 29°C. DMSO (Merck SA, Darmstadt, Germany, D8418) was used as a dissolvent for the AUTEN-67 AOBiome, Cambridge, MA, USA, AOB33340, T0501-7132) and AUTEN-99 AOBiome, Cambridge, MA, USA, AOB8904, T0512-8758) as well. Therefore, the control animals treated with the same amount of DMSO. To treat the animals, the stock solution was diluted to 100 μM in yeast suspension. All the treated animals were given fresh food every other day.

### Fluorescence microscopy

For microscopy the CO_2_-anesthetized animals were dissected and fixed in 4% of formaldehyde containing PBS solution for 20 minutes in room temperature, then washed with PBS (3×10 mins). 50ug/ml Hoechst nucleus stain in PBS: Glycerol (1:4) was used to cover the samples. The images were taken with a Zeiss inverted LSM800 confocal microscope (Hungary, Budapest, Institute of Biology at the ELTE University).

### Primary Mouse Hippocampal Cultures

The animal facility where the wild-type CD1 mice were housed maintained a temperature of 22 ± 1 °C and 12 h light/dark cycles, with ad libitum access to food and water. All experiments followed the local laws and standards for using animals in experiments, as well as the Hungarian and European Union legalisation. Embryonic cell cultures were derived from CD1 mice on embryonic days 17-18 and prepared as previously described ^65^. Cells were seeded onto poly-L-lysine-laminin-coated glass coverslips in 24-well plates. The medium contained NeuroBasal PLUS (ThermoFisher Scientific, Waltham, MA, USA, A35829-01) and the following supplements: 2% B27 PLUS (ThermoFisher Scientific, Waltham, MA, USA, A3582801), 5% FBS (PAN Biotech, Aidenbach, Germany, P30-3309), 0.5 mM GlutaMAX (ThermoFisher Scientific,Waltham, MA, USA, 35050-038), 40 μg/mL gentamicin (Merck SA, Darmstadt, Germany; G1397), and 2.5 μg/mL amphotericin B (Merck SA, Darmstadt, Germany; 15290-026). Cultures were maintained in vitro for 11–12 days at 37 °C with 5% CO_2_. On days 5 and 9, one-third of the medium was replaced with BrainPhys (StemCell Technologies, Vancouver, BC, Canada, 05790), with the addition of 2% SM1 (StemCell Technologies, Vancouver, BC, Canada, 05711), 40 μg/mL gentamicin, and 2.5 μg/mL amphotericin B. On the 11^th^ or 12^th^ day after plating, 10 μM AUTEN67 and/or AUTEN99 (Merck SA, Darmstadt, Germany, T0501-7132; Merck SA, Darmstadt, Germany, T0512-8758) were applied into the culture medium. In some cases, 50 nM Bafilomycin A1 (Merck SA, Darmstadt, Germany; B1793) was also used with the AUTENs. Twenty-four hours later, the cultures were fixed with 4% paraformaldehyde in PBS for 20 minutes.

### Immunostaining and Quantitative Microscopy in Fixed Hippocampal Cultures

Immunostaining of the cultures was performed using a protocol previously described ^66^ using anti-LAMP1 (mouse, 1:100; #1D4B; DSHB),, and anti-p62/SQSTM1 (Merck SA, Darmstadt, Germany rabbit, 1:2000, P0067) as primary antibodies and anti-mouse Atto 550 (Merck SA, Darmstadt, Germany, 1:500; A21237), and anti-rabbit Alexa Fluor 633 (Merck SA, Darmstadt, Germany, 1:500; #A21070) as secondary ones.

### Western Blot and Protein Isolation

We acquired samples from 3-day-old, 7-day-old, 14-day-old and 21-day-old adult female flies, kept at 29°C. We used our laboratory’s protocol for sample preparation ^26,67^. And the measurements were repeated multiple times. Our primary antibodies were anti-GFP (rat, 1:2500, Developmental Studies Hybridoma Bank, DSHB-GFP-1D2), anti-Tub84B (Merck SA, Darmstadt, Germany, mouse, 1:1000, T6199 [used as an internal control]). Secondary antibodies included anti-rat IgG alkaline phosphatase (1:1000, Merck Life Science, Darmstadt, Germany, A8438) and anti-mouse IgG alkaline phosphatase (Merck SA, Darmstadt, Germany, 1:1000, A8438). For label development the NBT-BCIP solution (Merck SA, Darmstadt, Germany, 72091) was dissolved in an AP buffer. ImageJ (1.54p) program was used for the densitometric analysis. In order to simultaneously determine the changes of the levels of free (degraded) mCherry-GFP and GFP, the data were normalized to that of αTub84B internal control. The quantities were adjusted to account for variations in the anti-Tub84B signals. We performed at least three repeats of the Western blot.

### Lifespan measurements

The flies, both control and treated, were placed on a new active medium every other day. Several parallel vials were used for each genotype and treatment type. Every day, the dead males and females were counted. A total of five parallel tubes containing 15 males and 15 females of each genotype and treatment were measured. The statistical evaluation was carried out using SPSS 17 software.

### Climbing

The CO_2_-anaesthetised were placed in a long thin glass tube (25 cm high, 1.5 cm in diameter). The animals were given 1.5 hours of regeneration time at 29°C before the assay. Gentle tapping triggered the negative geotaxis, causing the animals to fall to the bottom of the tube before climbing back up ^68^. Then the number of animals that reached the 6 cm line within 10 and 20 seconds and the 21.8 cm line within 20, 40 and 60 seconds was counted. Two parallel tubes were used for each treatment, each containing 15-15 animals. The male and female animals were measured separately. The assay was repeated three times, with a half-hour break between each repetition.

### Quantification and Statistical Analysis

We used RStudio 2023.12.1 for the analysis and to create the boxplots. For each dataset, we first performed a Shapiro–Wilk test to check for normal distribution. If the datasets were normally distributed, we tested the variance using an F-test. If the variances were equal, we compared the datasets using a two-sample t-test. If the variances were not equal, we used the Welch test. If one or both of the datasets to be compared had a non-normal distribution, we used the Wilcoxon rank-sum test (Mann–Whitney U-test) to compare the samples, which is equivalent to a two-sample t-test. We used boxplots to represent the evaluations, with the lower and upper edges of the boxes indicating the first and fourth quartiles. The thicker line in the middle represents the median. For fluorescent pictures, the dots represent individual samples. However, for the climbing assay, the dots show the percentage performance of animals tested at the same time. We have indicated the level of significance using stars: *** for p < 0.005, ** for p < 0.01, and * for p > 0.05.

## Ethics approval

This study was conducted under the approval of the Institutional Animal Ethics Committee of Eötvös Loránd University (approval number: PEI/001/1108-4/2013 and PEI/001/1109-4/2013). All methods were performed in accordance to the guidelines on research ethics of Eötvös Loránd for the use of experimental animals, in agreement with European Union and Hungarian legislation.

## Availability of data and materials

The datasets used and/or analysed during the current study are available from the corresponding author on reasonable request.

## Author Contributions

T.B. performed experiments on Drosophila samples, analysed data and wrote the manuscript. H.H. and E.I. performed experiments on Drosophila samples and analysed data. N.B. and K.S. carried out experiments on primary hippocampal neurons and analysed data and wrote the manuscript. T.K. designed experiments, analyzed data, and wrote the manuscript.

## Acknowledgments

Drosophila strains and reagents were kindly provided by Gábor Juhász (HUN-REN Biological Research Centre Szeged, Szeged, Hungary) and Huda Zoghbi (Baylor College of Medicine, Houston, TX 77030, USA). The authors also thank Regina Preisinger, Beatrix Supauer, Erzsébet Gatyás for the excellent technical assistance.

