## Supplementary material for "Investigation of autophagy-activating molecules in a glia-specific Spinocerebellar ataxia type 1 model": S.Figures

### Supplementary Figure 1

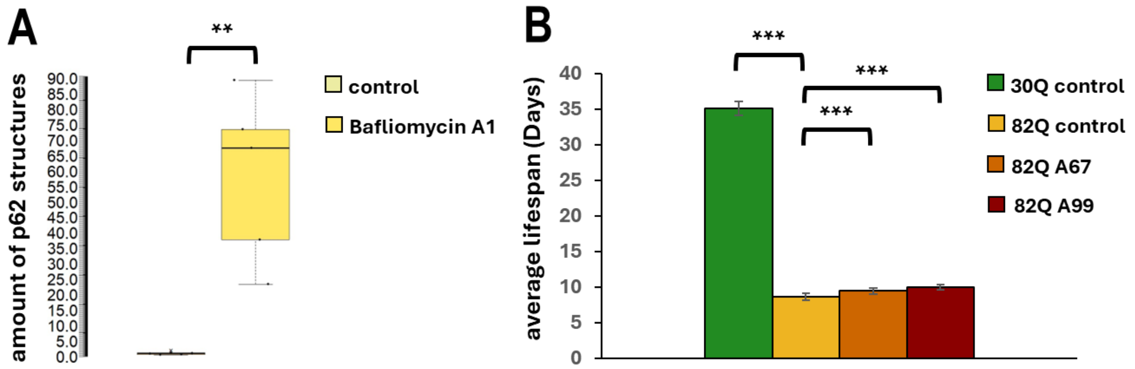


**S.Figure 1: Both AUTEN compounds significantly increase the mean lifespan of glia-specific SCA1 models. A)** The effects of AUTEN compounds were analysed in glial cells within mouse embryonic hippocampal primary cultures. Bafilomycin A1, an autophagy inhibitor that blocks autophagosome–lysosome fusion, was used as a negative control, and therefore an effect opposite to that of autophagy activators was expected. p62 is a substrate of autophagy, and its abundance is inversely correlated with the efficiency of lysosomal degradation. Bafilomycin A1 treatment resulted in a significant increase in the number of p62-positive structures. Representative images and quantitative analyses corresponding to this panel are shown in **Figure 2**. **B)** Using the glia-specific repo-Gal4 driver, wild-type (30Q) and mutant (82Q) human ATXN1 proteins were overexpressed in the glial cells of adult flies. The mean lifespan of 82Q animals was significantly reduced; however, this reduction was modestly but significantly rescued by AUTEN-67 (A67) and AUTEN-99 (A99) treatments. Kaplan–Meier survival curves corresponding to this analysis are shown in **Figure 3**.
